# Increased FOXL2 Expression Alters Uterine Structures and Functions

**DOI:** 10.1101/2020.03.06.981266

**Authors:** Rong Li, San-Pin Wu, Lecong Zhou, Barbara Nicol, John P. Lydon, Humphrey H-C Yao, Francesco J. DeMayo

**Author notes:** Summary Sentence: FOXL2 overexpression in the uterus induced epithelial stratification, blunted adenogenesis, increased fibrosis, and disrupted myometrium leading to impaired decidual responses and a similar transcriptome with human endometriosis. To whom correspondence should be addressed: Francesco J. DeMayo Ph.D., Chief, Reproductive and Developmental Biology Laboratory, National Institute of Environmental Health Sciences, 111 T. W. Alexander Dr. Research Triangle Park, NC 27709.

## Abstract

Transcription factor FOXL2 exhibits an increase in mRNA levels in eutopic endometrial biopsy in endometriosis patients. While FOXL2 is known of regulating sex differentiation and reproductive function, the impact of elevated FOXL2 expression on uterine physiology remains unknown. To answer this question, we generated mice with over expression of FOXL2 (FOXL2^OE^) in the female reproductive tract by crossing *Foxl2*^LsL/+^ with the *Pgr*^cre^ model. FOXL2^OE^ uterus showed severe morphological abnormality including abnormal epithelial stratification, blunted adenogenesis, increased endometrial fibrosis and disrupted myometrial morphology. In contrast, increasing FOXL2 levels specifically in uterine epithelium by crossing the *Foxl2*^LsL/+^ with the *Ltf*^icre^ mice resulted in the eFOXL2^OE^ mice with uterine epithelial stratification but without defects in endometrial fibrosis and adenogenesis, demonstrating a role of the endometrial stroma in the uterine abnormalities of the FOXL2^OE^ mice. Transcriptomic analysis of 12 weeks old *Pgr*^cre^ and FOXL2^OE^ uterus at diestrus stage showed a positive correlation of FOXL2^OE^ uterine transcriptome with human endometrium of endometriosis patients. Furthermore, we found FOXL2^OE^ mice were sterile. The infertility was caused in part by a disruption of the hypophyseal ovarian axis resulting in an anovulatory phenotype. The FOXL2^OE^ mice failed to show decidual responses during artificial decidualization in ovariectomized mice which demonstrates the uterine contribution to the infertility phenotype. These data supported that aberrantly increased FOXL2 expressions in the female reproductive tract can disrupt ovarian and uterine functions, particularly, may be involved in the progressions of endometriosis.

## Introduction

Endometriosis is a hormone dependent disease in which uterine endometrial cells grow outside the uterus. The most accepted hypothesis of the origins of endometriosis is that it is the peritoneal deposition of endometrial tissue from the retrograde menses that is not cleared by the body [1]. Endometriosis affects about 10 percent of women in the United States [2]. This disease results in pelvic inflammation, pain and infertility and is a significant women’s health issue. In women with endometriosis the eutopic endometrium and ectopic extrauterine tissues display altered gene expression signatures [3], including higher expression levels of Forkhead Box L2 (*FOXL2*) [4, 5]. Studying the role FOXL2 plays in regulating the biology of the reproductive tract will aid in understanding the cause of endometriosis and the development of treatments for this disease.

Forkhead Box L2 (FOXL2) is a transcription factor which contains a forkhead domain and a polyalanine tract. In addition to the human endometrium, FOXL2 is also expressed in the ovaries, the pituitary, including the gonadotrophs and some thyrotropes [4, 6, 7]. Germline mutation in FOXL2 has been associated with the blepharophimosis-ptosis-epicanthus inversus syndrome (BPES), characterized by eyelid malformation with or without primary ovarian insufficiency in humans [8]. Similarly, *Foxl2*^−/−^ mice display premature ovarian failure [9, 10]. Ablation of *Foxl2* at different stages of ovarian development demonstrated that *Foxl2* is a major regulator for the sex differentiation and maintenance of the ovary from the embryonic stage to adulthood [11–13]. In the pituitary, gonadotroph specific deletion of *Foxl2* in mice led to subfertility in both females and males due to Follicle-stimulating hormone (FSH) deficiency [14]. Both *in vitro* and *in vivo* models suggest that FOXL2-SMADs complexes can bind at the *Follicle Stimulating Hormone Subunit Beta* (*Fshb*) promoter to regulate its transcription [15–19]. FOXL2 also plays an important role in cancer development. It can suppress proliferation and promote apoptosis [20, 21]. Its mutation is present in most adult type granulosa-cell tumors and cervical cancers [20, 22, 23]. Taken together, FOXL2 is pivotal for the regulation of reproductive development and the hypophyseal-ovarian axis.

There are limited studies about FOXL2 functions in the uterus. FOXL2 is detected in the stroma and glandular epithelium of cow uterus throughout the estrous cycle with much higher levels during luteolysis, as progesterone treatment decreases its expression [24]. FOXL2 is also detected in the stroma and myometrium of mouse uterus, and conditional deletion of *Foxl2* by *Pgr*^cre^ reduced the stroma compartment while altering myometrial thickness and disrupting myometrial morphology [25]. In contrast, a recent study reported that FOXL2 is mainly detected in the epithelium and myometrium but not stroma in the mice [26]. In order to investigate the role of FOXL2 in embryo implantation, FOXL2 expression has been attenuated or enhanced in human endometrial cancer cell lines resulting in disruption of embryo attachment *in vitro* [26]. Despite of the contradictory expression patterns of FOXL2 in the uterus, these studies all suggested that FOXL2 may play a crucial role in endometrial homeostasis and function.

The goal of this work is to investigate the consequences of increased FOXL2 expression in the reproductive tract by employing the mouse model of uterine FOXL2 overexpression, *Pgr*^cre^*Foxl2*^LsL/+^ (FOXL2^OE^) and *Ltf*^icre^*Foxl2*^LsL/+^ (eFOXL2^OE^) mice. FOXL2^OE^ mice displayed multiple changes in uterine morphology, including impaired adenogenesis, altered uterine epithelial differentiation, increased collagen deposition and altered myometrial integrity, while eFOXL2^OE^ mouse uteris only showed epithelial stratification. Transcriptome of three months old FOXL2^OE^ uterus is positively correlated with human endometrium with endometriosis. FOXL2^OE^ mice were infertile with defective ovaries. The disrupted uterine functions were indicated by the abolishment of decidual responses upon artificial decidualization.

## Material and methods

### Mice

The *Rosa26-CAG-LSL-Foxl2* mice with FOXL2 overexpression (named as *Foxl2*^LsL/+^ in this paper), and *Pgr*^cre^ mice were described previously [13, 27]. *Ltf*^icre^ mice were kindly provided by Dr. Sudhansu K. Dey [28]. *Foxl2*^LsL/+^ mice were crossed with *Pgr*^cre^ mice to generate *Pgr*^cre^*Foxl2*^LsL/+^ mice, FOXL2 overexpression in female reproductive tract (FOXL2^OE^). To generate uterine epithelial FOXL2 overexpression mice, eFOXL2^OE^, *Foxl2*^LsL/+^ mice were crossed with *Ltf*^icre^ mice. All the mice were maintained on 129Sv and C57BL/6J backgrounds. All animal studies were conducted in accordance with the Guide for the Care and Use of Laboratory Animals published by the National Institutes of Health and animal study protocols approved by the Institutional Animal Care and Use Committee (IACUC) at the National Institute of Environmental Health and Sciences.

### Breeding Trial

8 weeks old *Pgr*^cre^ control mice and FOXL2^OE^ mice (N=6 for each genotype) were mated with stud males for 6 months. The copulation plug and delivery date for the first generation were recorded, and the pups delivered and survival rate were checked daily.

### Tissue collection

Postnatal Day (PND) 21, 3 months old (3M), 8 months old (8M) Pgr^cre^ and FOXL2^OE^ mouse ovary and uterus were collected at diestrus stage (N=6). The samples were fixed in 4% PFA, dehydrated, cleared and embedded in paraffin for histology and immunohistochemistry. The blood was collected from 3M diestrus *Pgr*^cre^ and FOXL2^OE^ mice for serum hormone analysis. 3M *Pgr*^cre^ (N=3) and FOXL2^OE^ (N=4) diestrus uterus were used for RNA-seq (N=3-4). 3M *Ltf^i^*^cre^ and eFOXL2^OE^ females were mated with stud males. The morning the plug was detected was defined as pregnancy day 0.5. The *Ltf*^icre^ and eFOXL2^OE^ mice (N=6) were sacrificed at pregnancy day 3.5, the uterus were collected and embedded in the paraffin.

### Superovulation

PND21 *Pgr*^cre^ and FOXL2^OE^ mice were injected with 3.25 IU of equine chorionic gonadotropin (eCG). 48h later, the mice were injected with 2 IU human chorionic gonadotropin (hCG). After 16h, the mice were sacrificed. The blood was collected for the serum hormone analysis. The oviducts were flushed for oocyte counting. The ovaries were fixed for paraffin embedding. N=6.

### Serum hormone analysis

The blood was collected by retroorbital bleeding. After clotting at room temperature (RT) for 1h, the blood samples were centrifuged at 2000g at RT for 10min. The supernatant was collected into a new tube and stored at −80ºC. The serum samples were shipped to The Center for Research in Reproduction Ligand Assay and Analysis Core, University of Virginia for luteinizing hormone, Follicle-stimulating hormone, estradiol and progesterone analysis. N=6.

### Artificial decidualization

*Pgr*^cre^ and FOXL2^OE^ mice were ovariectomized at 6 weeks old. After 2 weeks, the mice were subcutaneously (s.c.) injected with 100ng 17β-estradiol (E8875, Sigma) each day for three consecutive days, rested for two days, then given daily s.c. injections of 1mg progesterone (P0130, Sigma) and 6.7ng 17β-estradiol for three days. On the third day, 50ul corn oil was injected into one uterine horn 6h after the hormone injections. The mice were maintained with daily s.c. injections of 1mg progesterone and 6.7ng 17β-estradiol for five days, and the uteri were collected at the 6^th^ day. The uterine horns of both the oil injected and uninjected side were weighed. N=6.

### Histology and Masson’s trichrome staining

Paraffin embedded tissues were sectioned at 5 μm and a subset of sections were stained with hematoxylin solution, Harris modified (Sigma-Aldrich) and eosin (Sigma-Aldrich) for histology. A subset of sections was submitted to NIEHS histology Core for Masson’s trichrome staining. N=3.

### Immunohistochemistry and immunofluorescence

5μM paraffin sections were dewaxed and rehydrated for immunohistochemistry. After antigen retrieval, endogenous peroxidase blocking and serum blocking, they were incubated with primary antibody at 4ºC overnight, including FOXL2 (1:600, ab5096, abcam), HIS tag (1:300, ab9108, abcam), ESR1 (1:100, ACA054C, Biocare Medical), PGR (1:400, 8257, Cell signaling), FOXA2 (1:400, 8186, Cell signaling), P63 (1:800, 39692 Cell signaling), KI67(1:1000 ab15580 Abcam).

For immunohistochemistry, on the second day, the slides were incubated with 1:500 biotin-conjugated ant-rabbit (BA-1000 Vector laboratories), anti-goat (BA-9500 Vector laboratories) secondary antibody for 1h at room temperature respectively, followed by the ABC reagent (PK-6100 Vector laboratories) for 1h at room temperature. Signal was developed by DAB (SK-4105 Vector laboratories) for 30s. The slides were counterstained with hematoxylin, dehydrated, cleared and mounted by Permount medium (Thermo Fisher, Waltham, MA USA). The images were taken using Axiocam microscope camera (Zeiss). N=3.

For immunofluorescence, on the second day, the slides were incubated with Alexa Fluor® 647 goat anti-rabbit secondary antibody (1:300, ab150079, Abcam) for 1h at RT. The slides were mounted by ECTASHIELD® Antifade Mounting Medium with DAPI (H1200, Vector laboratories). The images were taken under Zeiss 710 confocal microscopy.

### RNA-seq analysis

Total RNA was isolated from 3 months old *Pgr*^cre^ and FOXL2^OE^ diestrus uteri using RNeasy mini kit (Qiagen). The library was prepared using TruSeq RNA Library Prep kit (Illumina) and subsequently sequenced using Nextseq 500. The sequencing reads with Quality score <20 were filtered using a custom perl script. The adaptor sequence was removed using cutadapt (v1.12). The reads were aligned to mm10 genome using STAR aligner (v2.5.2b) and counted using featureCounts (v1.5.0-p1) function in Subread program. The differential expressed genes (DEG) between *Pgr*^cre^ and FOXL2^OE^ were identfied using R package “DESeq2”. The threshold was set as “maximal FPKM>=1, unadjusted p<0.05, fold change >=1.5 (up-regulated) or =<−1.5 (down-regulated)”. The RNAseq data is deposited to NCBI Gene Expression Omnibus repository (GEO accession number) GSE140047.

The functions of the DEG were analyzed by Ingenuity Pathway Analysis (IPA, Qiagen) and DAVID Functional Annotation Bioinformatics Microarray Analysis [29, 30].

### Statistical analysis

The normality of the data was tested by Kolmogorov–Smirnov test. The equal variance of the data was tested by Levene’s test. The total number of pups, the number of ovulated oocytes, the serum levels of progesterone, 17β-estradiol, FSH and LH, the number of uterine glands, and the uterine gland penetration were compared by two tail, student’s t test. The significance was set at p<0.05.

## Results

### Increased endometrial FOXL2 expression in a subset of patients with endometriosis

Molecular profiles of eutopic endometrial tissues from 34 healthy human subjects and 28 patients who had minimal or mild endometriosis were examined to determine the relative expression level of FOXL2 between the two cohorts [5]. Normalized expression values of FOXL2, detected by the probe set 220102_at, were extracted directly from the series matrix file for comparison (GSE51981). The endometriosis specimens of this cohort exhibit higher levels of *FOXL2* expression compared with healthy endometrial biopsies (Suppl. figure 1A). Notably, in another cohort, FOXL2 levels are comparable in endometrial tissues between healthy and endometriosis subjects, but increased in the ectopic endometriosis tissues [4], suggesting that the elevation of FOXL2 expression occurs in the ectopic and/or eutopic endometriosis tissues. Based on these findings, we generated uterine *Foxl2* overexpression mice to model the observation on human and investigate the impact of increased FOXL2 expression in the uterus.

### Foxl2 transgene expression in the mouse uterus

Increased Foxl2 expression was achieved by crossing mice with a conditionally active transgene, *Foxl2*^LsL/+^ [13] with the *Pgr*^cre^ allele [27] generating the *Pgr*^cre/+^*Foxl2*^LsL/+^ (FOXL2^OE^) mouse. Immunohistochemical analysis was used to detect the expression of endogenous FOXL2 and the *Foxl2* transgene in the mouse uterus. In wild type mice,FOXL2 expression was observed in the endometrial stroma and blood vessels, and to a lesser extend in the epithelial and myometrial compartment of the uterus (Suppl. Fig. 1B-D) as previously reported [25]. The analysis of the uterine expression of the *Foxl2* transgene in the FOXL2^OE^, showed an increased expression in a subset of uterine luminal and glandular epithelium, and sporadic staining in the myometrium (Suppl. Fig. 1E-G). Due to the fact that FOXL2 is already highly expressed in the endometrial stroma cells of the *Pgr*^cre^ mice, it is difficult to distinguish the endogenous FOXL2 from the expression of the *Foxl2* transgene. Since the HIS Tag epitope was incorporated into the *Foxl2* transgene, the expression of the transgene in the endometrial stroma was determined by immunofluorescence staining for His tag [13]. Compared to the *Pgr*^cre^ mouse uterus (Suppl. Fig, 1 H-J), HIS tag displayed strong nuclear staining of the FOXL2 transgene in all compartments of the uterus (Suppl. Fig. 1K-M). This analysis demonstrated that the *Foxl2* transgene was expressed in not only the epithelial and myometrial compartment but also the stroma.

### Abnormal uterine morphology by Uterine FOXL2 overexpression

Macroscopic examination of the uterus showed that at PND21 the uterine size of the *Pgr*^cre^ and FOXL2^OE^ were comparable (Fig. 1A, D). However, the uteri of FOXL2^OE^ mice was thinner compared to the control mice at 3 and 8M at diestrus stage (Fig. 1B, C, E, F). Histological analysis identified alterations in all compartments of the uterus. Analysis of the uterine epithelium showed altered uterine epithelial cell differentiation. p63 is a marker for epithelial stratification [29]. While the control *Pgr*^cre^ mice showed a uterine epithelium lined with a single layer of columnal cells (Fig. 1G-I), the FOXL2^OE^ showed the presence of basal cells with P63 positive staining in glandular and luminal epithelium (Fig 1J-O). In addition to the altered uterine epithelial differentiation there was altered uterine gland morphology indicating a defect in adenogenesis.

**Fig. 1.**
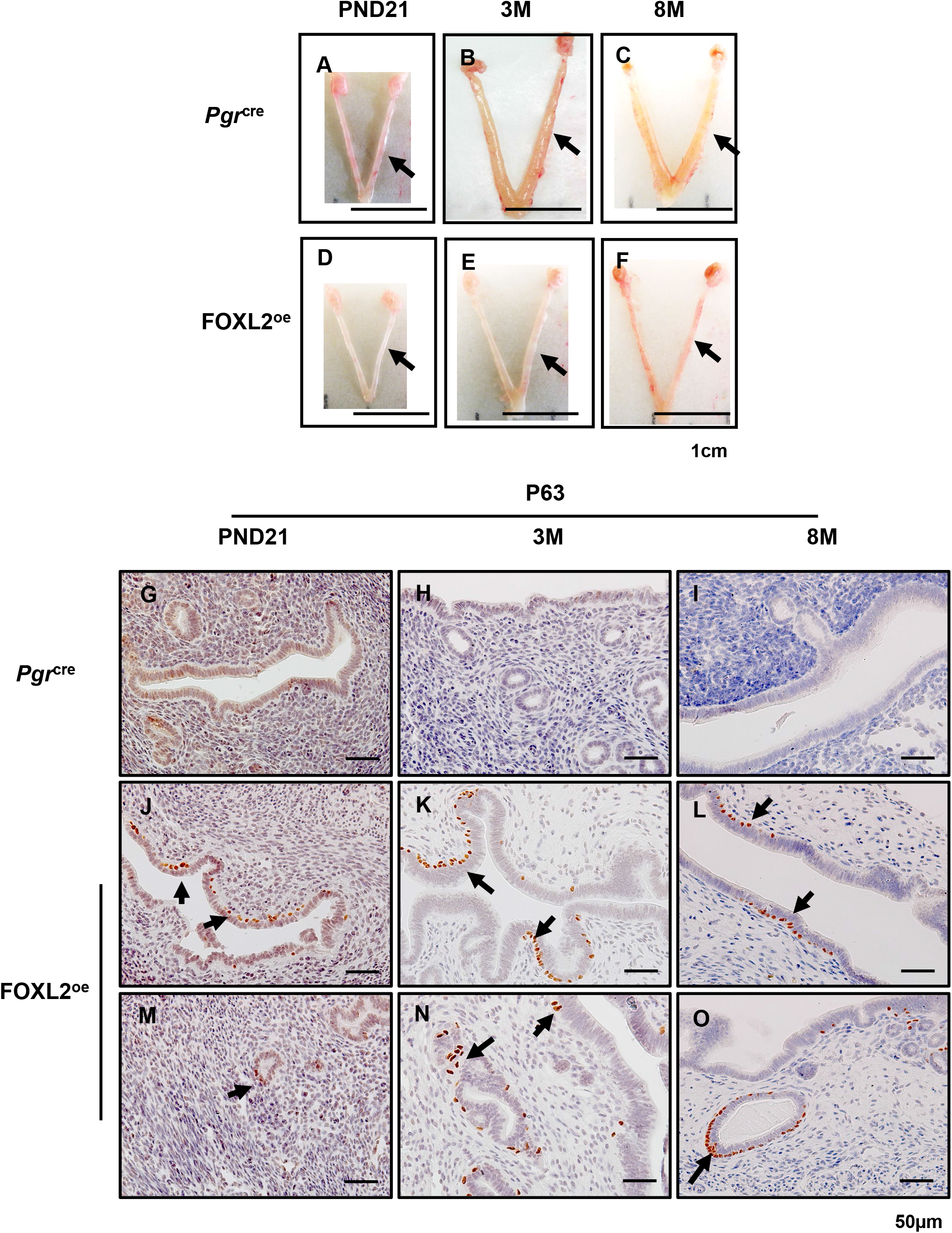
FOXL2^oe^ mice displayed thin uterus and stratified uterine epithelium. The representative uterine images of *Pgr*^cre^ (A-C) and FOXL2^oe^ mice (D-F) at PND21 (A, D), 3M (B, E), and 8M (C, F). The uteri were much thinner in FOXL2^oe^ mice compared to *Pgr*^cre^ mice at 3 and 6M. Basal cell marker, P63, staining in the uterus of *Pgr*^cre^ (G-I) and FOXL2^oe^ mice (J-O) at PND21 (G, J, M), 3M (H, K, N), and 6M (I, L, O). P63 positive basal cells in the uterus indicates epithelium stratification. They were found at the basal side of some luminal epithelium (J-L) and some glandular cells (M-O) in the FOXL2^oe^ uterus. PND: postnatal day; M: months old. Arrow indicates stratified epithelium. *p<0.05. N=3 for each genotype and age group.

Adenogenesis in rodents is a hormone independent process prior to puberty but is maintained by estrogen after puberty [31]. In the FOXL2^OE^ uterus, pre-pubertal gland development was already impaired at PND21, demonstrated by a decrease of the gland number and by the decreased penetration of the glands into the stroma (Fig, 2A, D, G, J, K). The defect in adenogenesis became more pronounced during the post-pubertal period. The *Pgr*^cre^ mice showed a remarkable increase in both the number of glands and gland penetration into the stroma compared to PND21. In contrast, the uterine glands of the FOXL2^OE^ remained at a lower number and few of them were able to penetrate the stroma. The impaired adenogenesis may result from a defect in gland growth or from the uterine stroma not providing the appropriate milieu for gland development. (Fig. 2B, C, E, F, H-K).

**Fig. 2.**
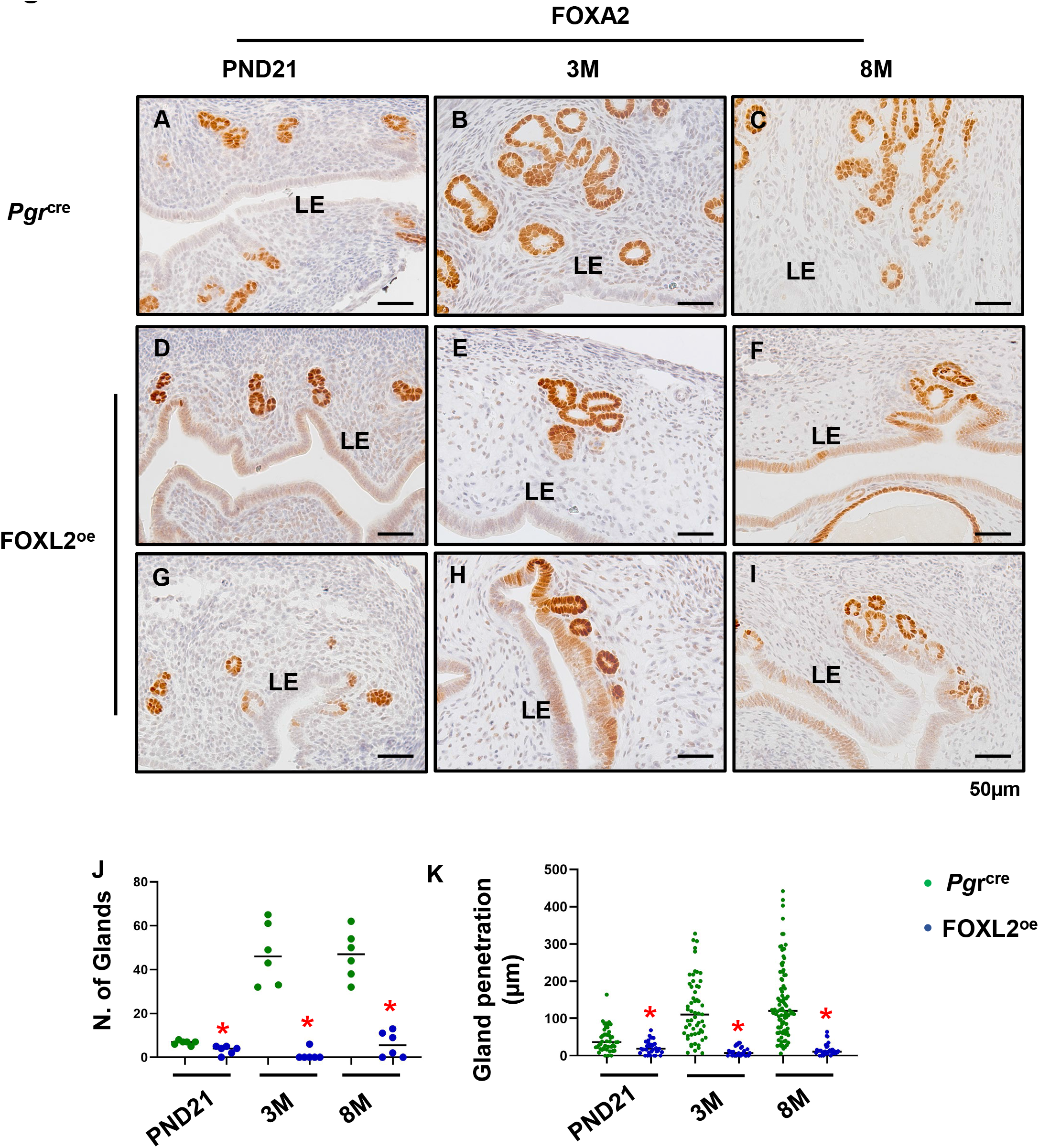
Disrupted adenogenesis in FOXL2^oe^ mice. FOXA2 labeled glands in *Pgr*^cre^ (A-C) and FOXL2^oe^ (D-I) uterus at PND21 (A, D, G), 3M (B, E, H) and 8M (C, F, I). The number of glands were calculated in two cross sections per mouse, in total 3 mice (J). The penetration of glands was defined as the closest distance of the glands to the adjacent luminal epithelium, and calculated in one longitudinal section per mouse, in total 3 mice (K). PND: postnatal day; M: months old. *p<0.05.

In order to determine the impact on FOXL2 expression on the endometrial stroma biology, the composition of the stromal extracellular matrix of the mouse uterus, *Pgr*^cre^ and FOXL2^OE^ mouse uteri was examined by Masson’s trichrome staining. Blue staining indicating collage deposition was increased with age in both *Pgr*^cre^ and FOXL2^OE^ mice (Fig. 3A-F). But at each age, the blue staining was much stronger in FOXL2^OE^ uterus suggesting increased collagen deposition (Fig 3A-F). The changes in the composition of the extracellular matrix may impede the ability of the glands to fully develop.

**Fig. 3.**
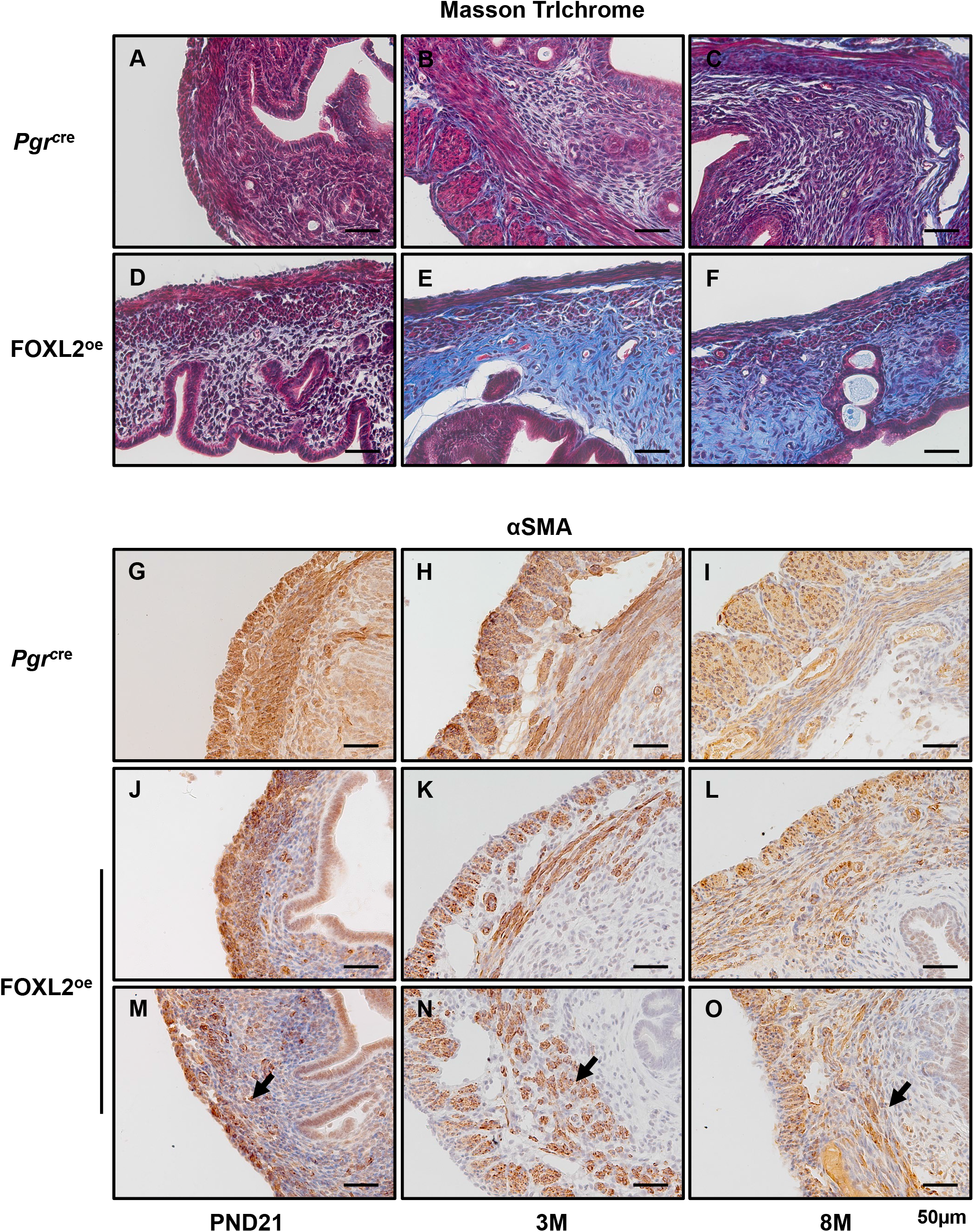
Stroma fibrosis and defective myometrium in FOXL2^oe^ mice. Masson’s trichrome staining in *Pgr*^cre^ (A-C) and FOXL2^oe^ (D-F) uterus at PND21 (A, D), 3M (B, E), 8M (C, F). Increased blue staining in the stroma of FOXL2^oe^ uterus suggested increased collagen deposition. αSMA staining in *Pgr*^cre^ (G-I) and FOXL2^oe^ (J-O) uterus at PND21 (G, J, M), 3M (H, K, N), 8M (I, L, O). The thickness of muscle layers was reduced and the inner muscle layer was discontinued in the FOXL2^oe^ uterus. PND: postnatal day; M: months old. Arrow indicates the discontinued muscle layer. N=3 for each genotype and age group.

In addition to the impact of *Foxl2* transgene expression on the uterine endometrium the transgene also affected the morphology of the myometrium. Immunohistochemistry was conducted to assay the expression of alpha smooth muscle actin (α-SMA), one of the common markers of mature myometrium [8]. The α-SMA positive myometrial cells were identified in both *Pgr*^cre^ and FOXL2^OE^ mouse uterus with no obvious changes of its staining intensity (Fig. 3G-O). However, while the myometrium of *Pgr*^cre^ mouse uteri showed the inner layer of myometrium to be a continuous layer of circular smoother muscle surrounding the endometrium (Fig. 3G-I), the inner smooth muscle layer of the FOXL2^OE^ mouse myometrium was discontinued at several loci (Fig. 3J-O, arrows). This abnormality was detected in the FOXL2^OE^ not the *Pgr*^cre^ uterus as early as PND21 (Fig. 3G, J, M), and remained at 3M (Fig, 3H, K, N) and 8M (Fig. 3I, L, O). The outer longitudinal smooth muscle layer increased proportionally with age in control mice from PND21 to 8M (Fig. 3G-I). The increase is dramatically diminished in the FOXL2^OE^ mice, resulted in hypotrophic muscle layer at 8M (Fig. 3L, O).

### Altered uterine epithelial differentiation but not adenogenesis, stromal fibrosis or myometrial structures by Epithelial FOXL2 overexpression

The FOXL2^OE^ mouse uterus exhibited FOXL2 overexpression in all the uterine compartments. In order to investigate the impact of altered FOXL2 functions specifically in the uterine epithelium, we bred the FOXL2^LSL^ mice with the uterine epithelial *Ltf^i^*^cre^ to generate epithelial overexpressing FOXL2, eFOXL2^OE^, mice. *Ltf^i^*^ce^ expresses Cre recombinase specifically in the uterine epithelium [28], FOXL2 and HIS-tag staining confirmed the FOXL2 overexpression is limited in a subset of luminal and glandular epithelium of eFOXL2^OE^ uterus (Suppl. Fig. 2).

Uterine morphology was assayed at D3.5 of pregnancy. The gross uterine morphology was similar between *Ltf*^icre^ and eFOXL2^OE^ females (Fig. 4A, B). The thread like uterus observed in FOXL2^OE^ mice (Fig. 1D-F) was not found in eFOXL2^OE^ mice. Histological analysis of the uterus of the eFOXL2^OE^ mice showed epithelial stratification with the presence of P63 positive basal cells, which was also observed in the FOXL2^OE^ mouse uterus (Fig. 4C, D). Unlike the FOXL2^OE^, eFOXL2^OE^ mouse uteri did not show any alteration in the number of uterine glands, as determined by FOXA2 staining (Fig. 4E, F), or in the uterine stromal fibrosis, as determined by Masson’s trichrome (Fig. 4G, H); or in their myometrium morphology, as determined by αSMA staining (Fig. 4I, J). These suggested that epithelial overexpression of FOXL2 impacted uterine epithelial stratification, but uterine gland development, stromal fibrosis and myometrial morphology were mainly affected by the extraepithelial overexpression of FOXL2. Based on this, FOXL2^OE^ mouse were used for further study of the impacts of FOXL2 overexpression in uterine transcriptome and functions.

**Fig. 4.**
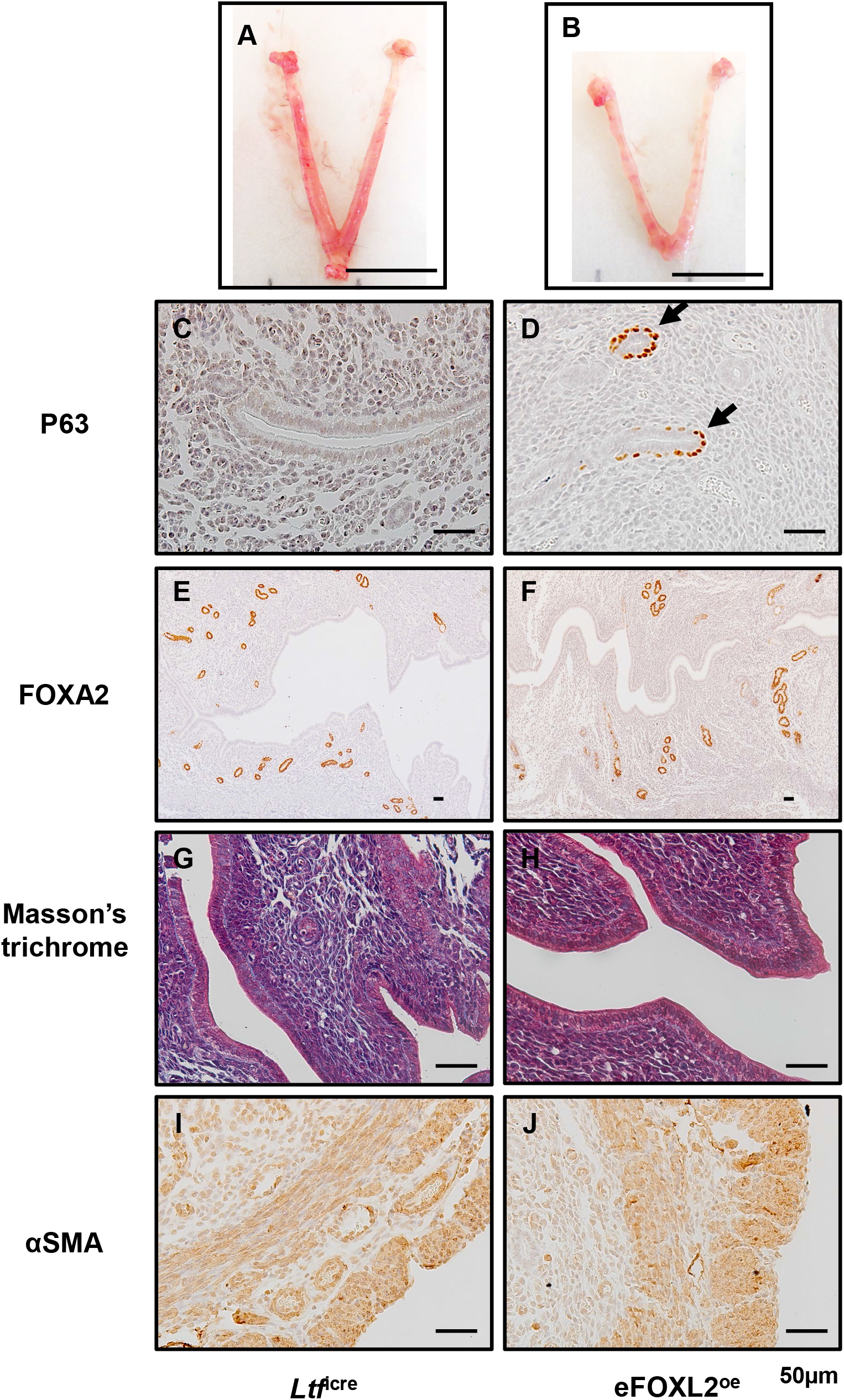
eFOXL2^oe^ exhibited epithelium stratification but no adenogenesis, stroma fibrosis and myometrial defects. The representative uterine images of *Ltf^i^*^cre^ (A) and eFOXL2^oe^ mice (B) at Pregnant D3.5. P63 (C, D), FOXA2 (E, F), Masson’s trichrome (G, H), and αSMA (I, J) staining in *Ltf^i^*^cre^ (C, E, G, I) and eFOXL2^oe^ (D, E, H, J) uterus. Basal cells with P63 positive staining was detected in the luminal and glandular epithelium of eFOXL2^oe^ uterus. No changes were observed in FOXA2 labeled uterine glands, Masson’s trichrome staining, and αSMA labeled muscle layers. Arrow indicates stratified epithelium. N=3.

### Foxl2 transgene impact on the uterine transcriptome

In order to determine the impact of the *Foxl2* transgene expression on the mouse uterus at the molecular level, RNA seq was conducted on the uteri of *Pgr*^Cre^ and FOXL2^OE^ of three months old. Since uterine transcriptome largely depends on the hormone regulation, we collected all the uteri at diestrus stage. In total, 3515 genes were differentially expressed in the FOXL2^OE^ compared to the *Pgr*^cre^ uterus (1746 upregulated and 1769 downregulated genes, respectively, Fig. 5A, the detailed gene list is in supplement excel file). Ingenuity pathway identified multiple signaling pathways altered in the FOXL2^OE^ uterus (Fig 5B, the full list of the altered pathways is in supplement excel file). The altered pathways included pathways regulating cell proliferation (cyclins and cell cycle regulation and estrogen-mediated S-phase entry, enhanced role of CHK proteins in cell cycle checkpoint control), extracellular matrix production (inhibition of matrix metalloproteases, suppressed collage receptor, GP6 and Hepatic Fibrosis/Hepatic Stellate activation), and Wnt/β-catenin signaling. These pathways likely contribute to the altered epithelial and stroma phenotype observed in the FOXL2^OE^ mouse uterus. Here we would like to mention that several collagen genes in the GP6 pathways were decreased in FOXL2^OE^ uterus such as *Col1a1*, *Col6a1*. In contrast, the collagen degradation genes in the inhibition of matrix metalloproteases were also suppressed in FOXL2^OE^ uterus, such as MMP2, MMP15. These results suggested that the inhibition of the matrix degradation may be the major reason for the increased collagen deposition in the FOXL2^OE^ uterine

The RNA-seq results identified several critical genes that are known to have an important role in the uterine phenotypes. We next validated the protein expression of selected genes. Since genes involved in cell cycle regulation were altered, *Ki67* expression was assayed. As expected, *Ki67* expression was decreased in our RNA-seq results (FC=−5.86, p<0.001), and immunohistochemistry confirmed the decreased protein levels of KI67 in both the epithelial and stromal compartments of the FOXL2^OE^ uterus (Fig. 6A, B). *Esr1* and *Pgr* are the major receptors for ovarian hormones and regulators of uterine function [32]. Our RNA-seq results showed mRNA levels of *Esr1* (FC=1.51, p<0.001) was increased and *Pgr* (FC=−1.26, p=0.035) was decreased in the FOXL2^OE^ uterus. Immunohistochemistry showed much higher expressions of ESR1 (Fig 6C, D, solid arrow) and much lower levels of PGR (Fig. 6E, F, open arrow) in the FOXL2^OE^ uterine epithelium. In the stroma, much more cells with weak ESR1 and PGR expression were found in the FOXL2^OE^ uterus compared to the *Pgr*^cre^ uterus (Fig. 6C-F, open arrow).

**Fig. 5.**
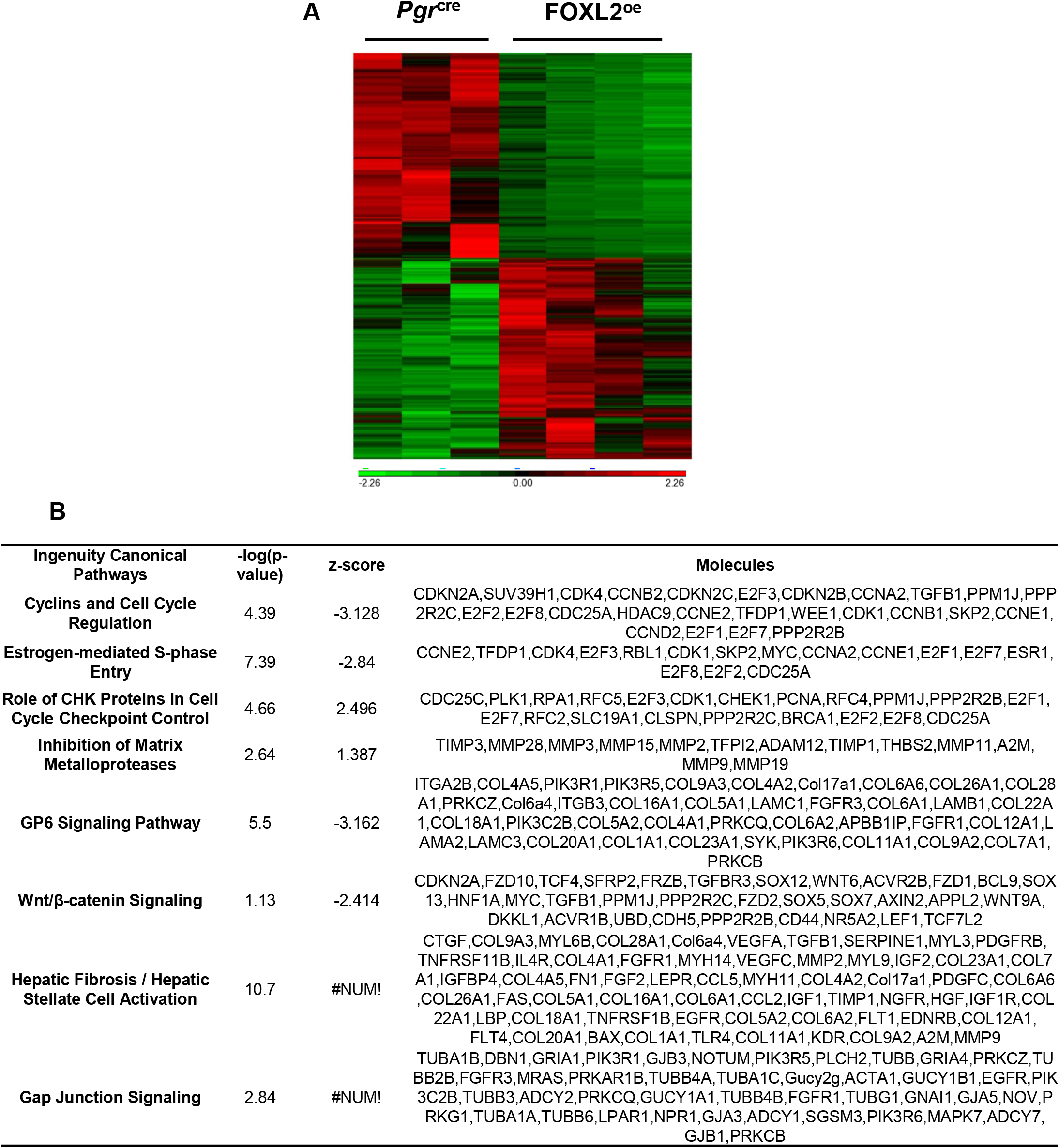
Transcriptomic changes of *Pgr*^cre^ and FOXL2^oe^ at 3M diestrus stage (p<0.05, fold change > 1.5, <−1.5). Heating map showed cluster of differentially expressed genes in *Pgr*^cre^ and FOXL2^oe^ (A). Ingenuity pathway analysis identified top altered signaling pathways (B). DAVID analysis indicates top altered cellular processes using up-regulated genes (C) and down-regulated genes (D). N=3-4.

**Fig. 6.**
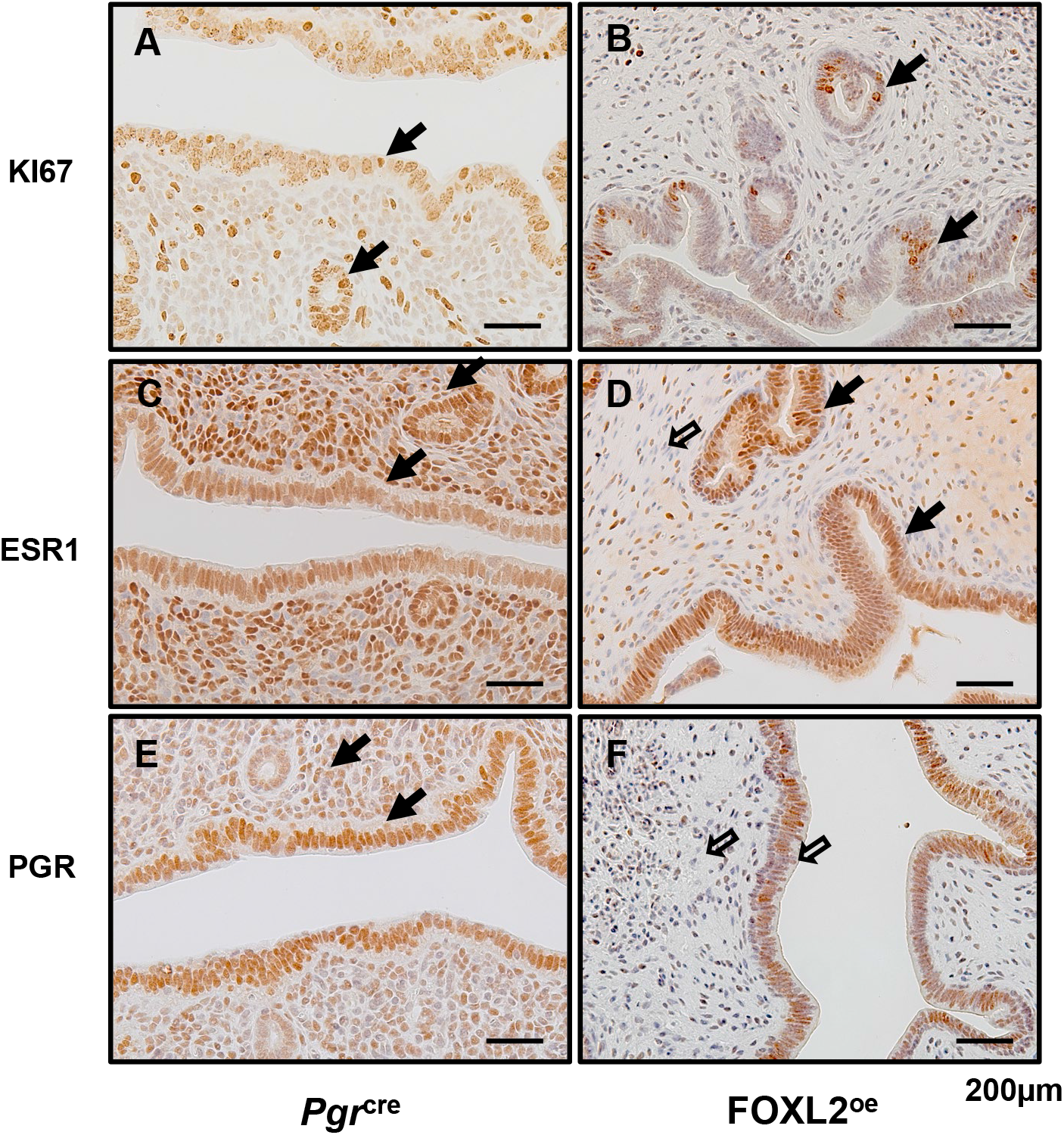
Protein expressions of KI67, ESR1 and PGR in FOXL2^oe^ uterus at 3M diestrus stage. Immunohistochemistry of KI67 (A, B), ESR1 (C, D), PGR (E, F) in *Pgr*^cre^ (A, C, E) and FOXL2^oe^ (B, D, F) uterus. In the *Pgr*^cre^ uterus, KI67, ESR1 and PGR was expressed in the nucleus of most epithelium and stroma cells. In the FOXL2^oe^ uterus, ESR1 was sporadically increased in the epithelium, PGR and KI67 was decreased in the epithelium of the FOXL2^oe^ uterus. Solid arrow indicates cells with strong staining. Open arrow indicates cells with weak staining. N=3.

As mentioned above, FOXL2 is increased in several human endometriosis tissues [4, 5]. To investigate the possible correlation of the FOXL2 mouse model with the human endometriosis, we used Nextbio [33] to determine if the FOXL2 transcriptome correlated with transcriptome of any human endometrial data sets. This analysis identified a significant positive correlation of the FOXL2^OE^ transcriptome with multiple human endometriosis transcriptome data sets (GSE7305, GSE5108, GSE87809, GSE51981, GSE4736, Fig. 7A, Suppl. Fig. 3A-D), including human endometrosis tissues compared to the normal normal endometrium and ectopic compared to eutopic endometriosis tissues [5, 34–37]. As expected, a positive correlation of the transcriptome was established in the dataset GSE51981 (Suppl. Fig 3C), in which the increased *Foxl2* mRNA levels were identified in the human endometrium with endometriosis compared to the normal human endometrium (Suppl. Fig. 1A). Among them, the most correlated data set was collected from the human ovarian endometriosis tissues that was compared to the control endometrium of the same patients using microarray analysis revealing 5864 up-regulated and 6361 down-regulated genes (Fig. 7A, [35]). We found 1816 genes were common between the two data sets. Of these, 1152 (63%) genes showed expression changes in the same direction. We analyzed the functions of the 1816 overlapping genes using IPA and DAVID and identified several common pathways between these two data sets (Fig 7B, C). These pathways were regulation of cell proliferation and death, cellular matrix, cell migration and adhesion, regulation of immune system process, vasculature development, and pathways that can be associated with epithelium differentiation problems such as tube and gland development. These are pathways which are known to be critical for the progression of endometriosis [38–40].

**Fig. 7.**
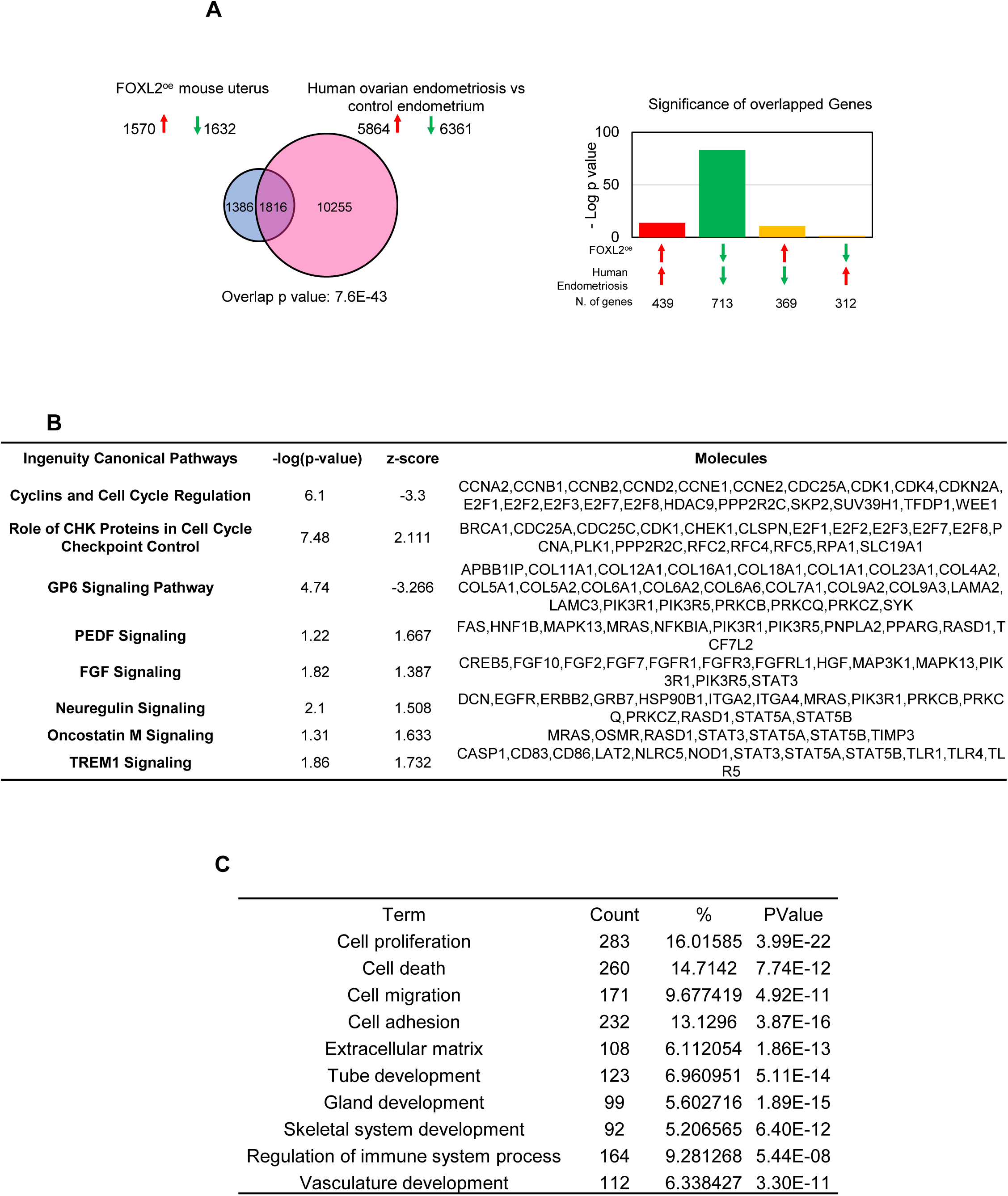
FOXL2^oe^ transcriptomic was positively correlated with human endometriosis. The most positively correlated human endometrium transcriptome determined by Nextbio was the comparison between ovarian endometriosis tissue with control endometrium in the same patient (GSE7305 [35], A). The number of up- and down-regulated genes were labeled accordingly near the up and down arrows. The number of overlapped genes between both datasets were plotted in the venn diagram. The significance of the overlap was showed below the venn diagram. Furthermore, the 1816 overlapping genes were divided into four groups based on its up- or down regulated in either dataset. The significance and number of overlapped genes in these four groups were displayed in the bar graph. The IPA (B) and DAVID (C) analysis of the 1816 overlapping genes between FOXL2^oe^ and human endometriosis identified multiple altered pathways that may relate with endometriosis.

### Infertility in FOXL2^OE^ mice

As we already observed multiple uterine functions in the FOXL2^OE^ mice, we decided to investigate the impact of the *Foxl2* transgene expression by a 6-month breeding trial. Adult *Pgr*^cre^ and FOXL2^OE^ female mice were placed with a male mouse and allowed to breed for 6 months. During the six months, *Pgr*^cre^ females produced 36±2.35 pups/ female, while FOXL2^OE^ females produced no offspring (Fig. 8A), This demonstrated that the FOXL2^OE^ female mice were infertile. In order to determine the cause of infertility, female mice were examined for the presence of a postcoital vaginal plug after being placed with a male mouse. No copulation plug was ever detected in FOXL2^OE^ females during the breeding trial. Since mating occurs in female mice during the estrus stage of the cycle, the ability of the FOXL2^OE^ mice to undergo a normal estrus cycle was analyzed by vaginal cytology over a two-week period. We found FOXL2^OE^ female mice did not show the presence of a vaginal estrus cycle and were in a constant state of diestrus (Fig 8B, C). Analysis of estrogen and progesterone levels in the FOXL2^OE^ mice reveals that they were comparable with those of *Pgr*^cre^ mice in the diestrus stage (Suppl. Fig. 5A, B). This confirmed the continuous diestrus stage in the FOXL2^OE^ mice not only reflected on the vagina cytology but also by the ovarian hormone levels.

**Fig. 8.**
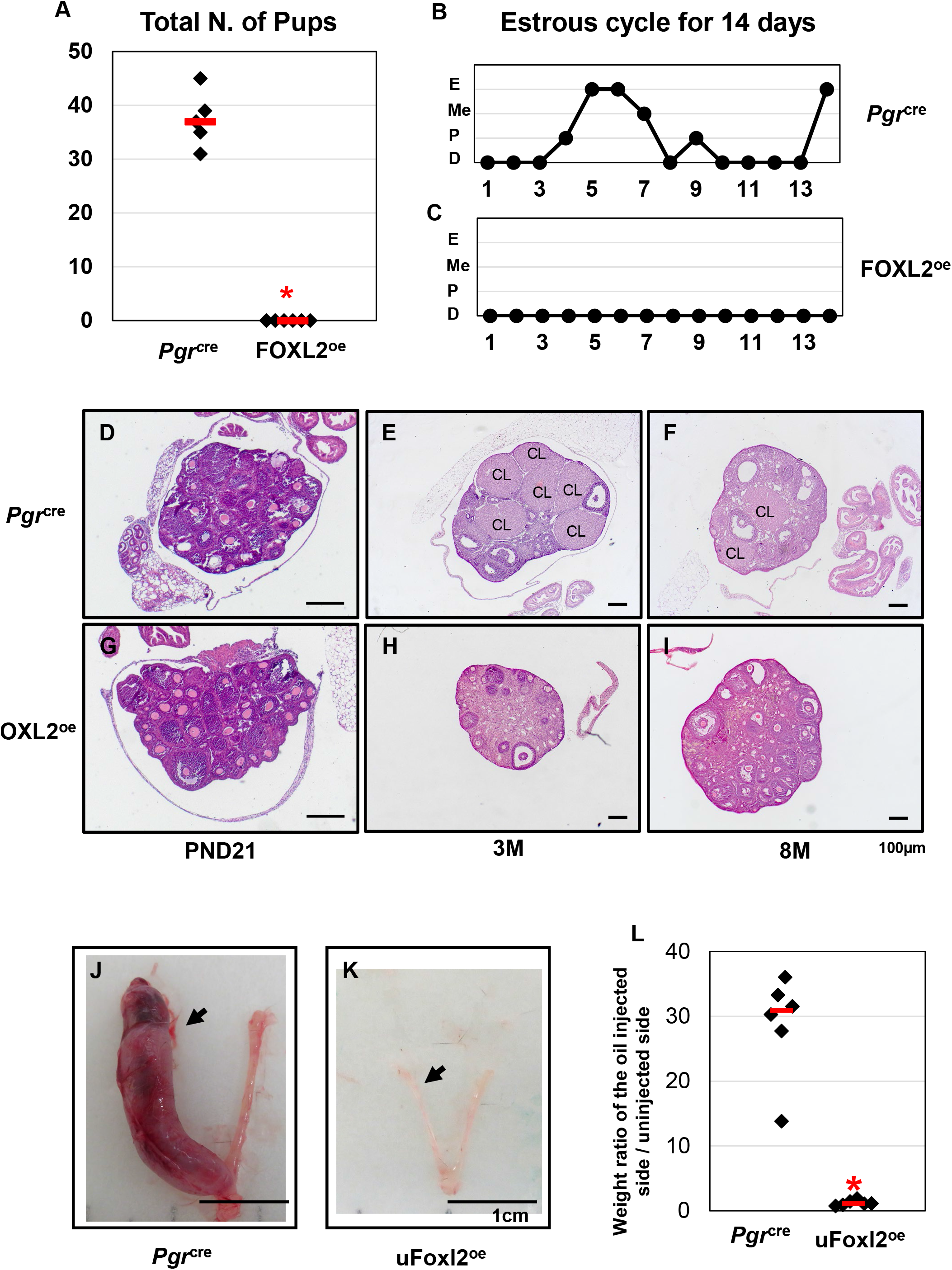
FOXL2^oe^ mice were infertile with abolished decidual responses. All the *Pgr*^cre^ mice but none of the FOXL2^oe^ mice delivered any pups during 6-month breeding trial (A). Representative pictures of estrous cycle showed continuous diestrus in FOXL2^oe^ mice (C) in contrast to the estrous cycle showed in *Pgr*^cre^ mice (B) during 14-day period. Representative ovarian histology pictures of *Pgr*^cre^ (D-F) and FOXL2^oe^ (G-I) virgin mice at diestrus stage at the age of PND21 (D, G), 3M (E, H) and 8M (F, I). FOXL2^oe^ ovaries were depleted of corpus luteum. The representative pictures of the hormone primed uterus after 5-day oil injections in the *Pgr*^cre^ (J) and FOXL2^oe^ (K) mice. The weight ratio of the oil injected side over the un-injected side uterine horn (L) confirmed the decidualization occurred in all the *Pgr*^cre^ but none of the FOXL2^oe^ mice. Arrow indicates the oil injected side. D: diestrus; P: proestrus; Me: metestrus; E: estrus; PND: postnatal day; M: months old; CL: corpus luteum. Arrow indicates oil injected uterine horn. n=6 for each genotype and age group.

### Altered ovarian function in FOXL2^OE^ mice

The estrus cycle of mice is driven by steroid hormones produced by the ovaries. Since *Pgr* is transiently expressed the ovarian corpus luteum [41], we examined the ovaries to determine whether the acyclicity was caused by any ovarian defects. At PND21, the ovaries contained most small and medium follicles in both *Pgr*^cre^ and FOXL2^OE^ mice (Fig. 8D, G). At 3M and 8M, the ovaries of *Pgr*^cre^ mice showed follicles at different stages and multiple corpora lutea (Fig. 8E, F). However, the FOXL2^OE^ mouse ovaries showed normal follicular development until antral follicle but complete absence of corpora lutea (Fig. 8H, I).

The lack of corpora lutea in the FOXL2^OE^ could be intrinsic to the ovary or could be due to a defect in neuroendocrine regulation of the ovulatory process. In order to determine the reason for the lack of corpora lutea in the FOXL2^OE^ mice, we first investigated the expression of the *Foxl2* transgene in these mice. At 3M, both *Pgr*^cre^ and FOXL2^OE^ ovaries from the mice in diestrus showed high levels of endogenous FOXL2 in the small and medium follicles (Suppl. Fig. 4A, B). However, His-Tag staining detected no strong nuclear staining in *Pgr*^cre^ and FOXL2^OE^ ovaries suggesting no *Foxl2* transgene expression in FOXL2^OE^ mouse ovaries (Suppl. Fig. 4C, D). Therefore, the ovarian phenotype may not be due to ovarian expression of the *Foxl2* transgene since the follicles could not develop to a stage where the *Pgr*^cre^ would activate the expression of the transgene.

We next examined if the ovarian phenotype could be rescued by administering a super ovulatory regimen of gonadotropins to mice. We found 7/10 *Pgr*^cre^ and 3/10 FOXL2^OE^ ovulated in response to the superovulation of gonadotropins. *Pgr*^cre^ mice that ovulated produced 34.3±4.5 oocytes and FOXL2^OE^ mice that ovulated produced 8.3±3.8 oocytes (Suppl. Fig. 4I). All mice showed corpora lutea in the ovary (Suppl. Fig. 4G, H, J), and *Foxl2* transgene expression was identified by His-tag in the peri-ovulatory follicles and the corpus luteum of the FOXL2^OE^ mice (Suppl. Fig. 4F, H). Although the ovulation capability was reduced in the FOXL2^OE^ mice, the formation of corpus luteum in the FOXL2^OE^ mice indicates that the corpus luteum can and does form in FOXL2^OE^ ovary under the appropriate hormone stimulation.

Since one major function of the corpus luteum is progesterone synthesis, we checked the serum progesterone levels of one litter of 21 days old mice after superovulation to test whether these corpus lutea were functional. Both the *Pgr*^cre^ and four FOXL2^OE^ mice displayed increased progesterone levels in comparison to the diestrus mice with no difference between the *Pgr*^cre^ and FOXL2^OE^ mice (Suppl Fig. 5A, E). This indicated that FOXL2 overexpressed corpus luteum capable of producing progesterone. Based on these results, we proposed that the cause of the FOXL2^OE^ ovarian dysfunctions was a defect in neuroendocrine control of ovulation. To evaluate the functions of the pituitary, the direct upstream regulator of ovary, we assayed the serum levels of two gonadotropins produced by the pituitary: Follicle stimulating hormones (FSH) and Luteinizing hormone (LH) showed no difference in serum levels (Suppl Fig. 5C, D). Although the pituitary is capable of production FSH and LH in the FOXL2^OE^ mice, we have not ruled out that the appropriate cyclic regulation of these gonadotropins is altered. This determination is beyond the focus of this report as the primary focus is the investigation of the phenotypic consequences of the *Foxl2* transgene on the mouse uterus.

### Impaired ability of the FOXL2^OE^ uterus to undergo a decidual reaction

It is true that the ovarian defects led to the infertility of the FOXL2^OE^ mice, but considering the multiple uterine morphological changes, it is possible that the uterine functions might also be altered. Since one major function of the uterus is to accommodate the embryo implantation by decidualization, the ability of the uterus to undergo a decidual reaction was assayed. The decidualization of the uterine stroma, which involves both stroma and epithelial signaling, is critical is to support pregnancy [42]. To avoid the ovary defects, female *Pgr*^Cre^ and FOXL2^OE^ mice were ovariectomized and administered an ovarian hormonal regimen combined with trauma to one uterine horn to assess the decidual response. The control mice showed an increase in the uterine size of the traumatized horn as compared to the untraumatized horn indicative of a decidual response (Fig. 8J, L). In contrast, the size of the traumatized horn of the FOXL2^OE^ mice did not increase at all indicating the complete absence of a decidual response (Fig. 8K, L). This demonstrates that altered FOXL2 expression impairs the hormonal differentiation of the uterus necessary to support pregnancy.

## Discussion

In this study we found FOXL2 overexpression in PGR expressing cells severally impaired uterine structures and functions. These mice showed altered uterine epithelial differentiation, disrupted adenogenesis, altered endometrial stroma extracellular matrix, impaired myometrium development. In contrast, the epithelial specific FOXL2 overexpression only led to epithelial stratification suggesting the importance of extraepithelial FOXL2 contribution in the uterine phenotypes. Since FOXL2 expression is increased in humans with endometriosis we compared the FOXL2^OE^ uterine transcriptome with the transcriptome of endometriosis patients and identified positive correlations between FOXL2^OE^ uterine transcriptome and multiple transcriptome of endometriosis patients. Additionally, the female FOXL2^OE^ mice were sterile with constant diestrus. The ovaries showed no signs of ovulation or corpora lutea formation which can be rescued by superovulation implying hypohpyseal gonadal axis defects. The uterus cannot respond to the decidual cue under the appropriate hormone priming patterns. All these results demonstrate a pleiotropic role of FOXL2 overexpression in female reproductive system.

Since we observed constant diestrus in FOXL2^OE^ mice in which stage estrogens and progesterone are at relatively low levels, it is natural to assume that these low levels of ovarian hormones might explain some of the FOXL2^OE^ uterine phenotypes. However, the uterus of the FOXL2^OE^ mouse does not phenocopy what is observed in the ovariectomized mouse models. First ovariectomy has never been positively correlated with epithelium stratification, instead, estrogen is required for vagina epithelium stratification [43]. The stratification was also observed in the mice with epithelial specific overexpression of FOXL2 indicating a phenotype of uterine origin. Second, neither ovariectomy nor ablation of PGR and ESR1 impair prepubertal adenogenesis as observed in the FOXL2^OE^ mouse [44–46]. Since ovariectomy does impair post-pubertal glandular development, the alteration in ovarian hormone production may be partly responsible for the reduction of adenogenesis at 3 and 8M. Third, ovariectomy has been reported to decrease collagen deposition in the uterus [47]. Finally, the reduced ovarian hormone might account for the hypotrophic myometrium but not the discontinued smooth muscle layer [48]. According to these evidences, we conclude that the FOXL2^OE^ uterine defects are of uterine origin.

The stratified squamous uterine epithelium has been observed in both FOXL2^OE^ and eFOXL2^OE^ uterus suggest epithelial *Foxl2* transgene expression is sufficient to induce the epithelial differentiation phenotypes. Previous studies reported multiple mouse models with enhanced estrogen signaling may stimulate the uterine stratification [49–52]. Considering the low estradiol scenario in this FOXL2^OE^ mice which are at continuous diestrus stage, alternative pathways might be involved. In our studies we found that the Wnt/β-catenin pathway was suppressed in the 3M FOXL2^OE^ mouse uterus. Wnt/β-catenin signaling interacts with multiple estrogen signaling [53, 54]. Mice with either *Pgr^cre/+^* induced deletion of β-catenin in *Pgr^cre/+^Ctnnb1^dd^* mice [55], or with global knockout of *Wnt7a* [56] mice exhibit epithelium stratification in the uterus. It is possible that FOXL2 can promote epithelial stratification through suppressing the Wnt/β-catenin pathway. The FOXL2 negative regulation on Wnt/β-catenin has also been reported in other studies. *Pgr^cre/+^* induced deletion of FOXL2 upregulates Wnt4 and Wnt7a in the mouse uterus at PND25 [25]. FOXL2 and β-catenin have been shown opposite functions on cell proliferation in ovarian cells [57].

Normal gland development in the eFOXL2^OE^ uterus suggested that epithelial FOXL2 overexpression does not impair adenoegenesis. For the pre-pubertal adenogenesis, it is possible that stroma FOXL2 exert its effects also through Wnt/β-catenin signaling, which has been identified to be important for the pre-pubertal adenogenesis [58–61]. For the post-pubertal adenogenesis, it is also true that that low levels of estradiol might inhibit the gland growth, but one unique phenotype in FOXL2^OE^ mice is the severely impaired penetration of the glands into the stroma. It is possible that without the appropriate penetration, the glands cannot elongate and branch further leading to a decrease in the number of glands. The increased collagen deposition may act as the physical obstacle of gland penetration, suggesting the importance of the endometrial stroma extracellular matrix in allowing normal uterine gland development.

Increased collagen deposition was also found in the *Pgr^cre^* induced Transforming Growth Factor Beta Receptor 1 (TGFBR1) or Smoothened (SMO) overexpression mouse models [62–64]. However, these studies did not exclude potential confounding factors from the ovary, so it is still unclear whether the uterine TGFBR1 and SMO are the major players. One study using FOXL2−/− male mice shows retarded cartilage and bone formation [65], suggesting a positive role for FOXL2 in regulation of cartilage formation. Our study presents a direct evidence showing that uterine FOXL2 overexpression may increase collagen deposition. The inhibition of matrix metalloprotease signaling is mainly composed of matrix degradation related genes. This signaling is enhanced in the 3M FOXL2^OE^ uterine transcriptome indicates that FOXL2 can also increase collagen through inhibition of collagen degradation.

A previous study also reported a disorganized smooth muscle layer in *Pgr^cre^* induced FOXL2 knockout mice [25], and together with our study, these observations indicate that appropriate FOXL2 levels are required for the myometrium development. Similarly, either deletion or overexpression of TGFBR1 leads to disrupted smooth muscle layers suggesting the subtle roles of TGFβ pathways [64, 66]. Additionally, overexpressed TGFBR1 also reduced pre-pubertal adenogenesis [58], Since the FOXL2-SMAD4 complexes acted synergistically in the pituitary [15, 19], it is possible that FOXL2 also collaborated with the TGFβ pathways in regulation of myometrium development.

Endometriosis is prevalent in women at reproductive age [2], but the pathologic mechanisms leading to the disease remain unclear. FOXL2 overexpression has been found in human endometriotic lesions [4]. Our FOXL2 overexpression mouse model further indicates a positive correlation with human endometriosis. Pathways including fibrosis, inflammation, cell adhesion and migration are among those that have been shown to be involved in regulating endometriosis progressions [38, 67], and were altered in the uterine transcriptome of the 3 months old FOXL2^OE^ mice. Immunohistochemistry also showed decreased progesterone receptor in the FOXL2^OE^ uterus. This is consistent with the protective role of progesterone in endometriosis [68], and the lower progesterone receptor in endometrotic lesions compared to the eutopic endometrium [69, 70]. This study has identified a FOXL2 signature that may contribute to the endometriotic phenotype observed in patients.

The defect in the estrus cycle and ovarian function in the FOXL2^OE^ is most likely due to a disruption of the hypohpyseal gonadal axis. The *Foxl2* transgene is detected in the ovary of the FOXL2^OE^ mouse, yet FOXL2^OE^ mice were able to respond to exogenous gonadotropins and ovulate and produce normal levels of progesterone. The presence of corpus luteum in the super ovulated ovary suggested that the FOXL2 overexpression in the ovary did not affect its ability to develop a corpus luteum. Although we could not detect any changes in normal circulating gonadotropins this could not rule out the impact of FOXL2 on pituitary function. Since deletion of FOXL2 has been reported to affect gonadotropin production by the pituitary leading to ovarian defects, the alteration of FOXL2 expression may disrupt the timing of gonadotropin release necessary to stimulate appropriate follicular development and ovulation [14, 71]. The precise effects of FOXL2 overexpression on pituitary development requires further investigation.

In addition to the structure alterations, the failure of the FOXL2^OE^ uterus to respond to the decidual cue indicates the functions of the uterus to support pregnancy were also impaired. Ovariectomy with exogenous hormone treatment eliminates the ovarian effects and indicates an intrinsic uterine defect. It is possible that the structure abnormality might lead to the function defects. The stratified epithelium may lose its ability to support the pregnancy. Previous studies identified several mutant mouse models with epithelium stratification were infertile, such as *Pgr^cre/+^Ctnnb1^dd^* mice [55], *Wnt7a−/−* [56] mice, and *Pgr^cre/+^Sox17^dd^* mice [49]. However, since these mice exhibited multiple phenotypes, it is still too presumptions to link the epithelial stratification with pregnancy failure. In contrast, the presence of functional glands is well known to be critical for uterine decidualization probably through LIF signaling [72–74]. The severely retarded adenogenesis in the FOXL2^OE^ uterus might be one of the major players in the failed decidualization. Besides, endometrial decidualization is promoted by mesenchymal-epithelial transition (MET) [75], while fibrosis is positively correlated with epithelial-mesenchymal-transition (EMT) [76]. Therefore, the exaggerated fibrosis observed in the FOXL2^OE^ uterine stroma might also blunt the decidualization through inhibiting the MET process.

Our study first reported that the *Pgr^cre^* induced *Foxl2* transgene expression can alter the uterine structures and functions, including adenogensis, epithelium stratification, collagen deposition, and myometrial development. It provides a model to investigate the role of the stroma extracellular matrix in uterine gland development. It also provides new evidence to positively correlate FOXL2 overexpression with endometriosis progression. Further studies are required to identify the underlying mechanism and the therapeutic potentials of FOXL2 in human.

## Acknowledgments

The authors acknowledge the Epigenomic and DNA Sequencing Core, The DNTP Clinical Pathology Core, the Integrative Bioinformatics Supportive Group, the Fluorescence Microscopy and Imaging Center and the Comparative Medicine Branch at NIEHS for their research support. The authors acknowledge Research in Reproduction Ligand Assay and Analysis Core, University of Virginia for serum analysis. The authors appreciated Dr. Tianyuan Wang’s consultation on bioinformatic analysis, Mr. Linwood Koonce for mouse colony management, Ms. Mita Ray for sharing the reagent, and Dr. Sylvia Hewitt for the constructive input on the paper and Ms. Janet DeMayo for the proof reading.

## Notes

Grant support: This work was funded by an Intramural Research Program of the National Institute of Health (NIH) Z1AES103311-01 to F.J.D. and a NIH / National Institute of Child Health and Human Development (NICHD) RO1 HD042311 to J. P. L.

