## Supplement Figures for "Increased FOXL2 Expression Alters Uterine Structures and Functions"

**Suppl. Fig. 1**. *FOXL2* expression was elevated in the human endometriosis tissues and FOXL2 was overexpressed in the FOXL2$^{oe}$ mouse uterus. *FOXL2* mRNA levels were increased in the endometriosis compared to the non-endoemtriosis endometrium in human (A). FOXL2 is mainly expressed in the stroma and muscle layers and much weaker in the epithelium in the *Pgr*$^{cre}$ mice (B-D). FOXL2 is detected in some luminal (E) and glandular (F) epithelium, and highly expressed in the muscle layer in FOXL2$^{oe}$ mice (G). Immunofluorescence of HIS-tag indicated FOXL2 overexpression is detected in the epithelial, stroma and some muscle cells in the FOXL2$^{oe}$ mouse uterus (K-M), in comparison with the relatively low and punctuate HIS staining in the *Pgr*$^{cre}$ mouse uterus (H-J). HIS-tag: red; DAPI: Blue. Arrow indicates cells with positive staining. N=6.

**Suppl. Fig. 2**. FOXL2 was overexpressed in the uterine epithelium of eFOXL2$^{oe}$ mice. Endogenous FOXL2 is detected in the stroma and much lower in the epithelium and myometrium in the Ltf$^{icre}$ uterus (A-C), Low levels of punctuate HIS-tag staining was observed in the cell nucleus of the Ltf$^{icre}$ uterus (G-I). FOXL2 (D-F) and HIS-tag (red staining, J-L) are highly detected in a subset of luminal epithelium (D, J) and glandular epithelium (E, K) of eFOXL2$^{oe}$ uterus, but not found in the stroma and myometrium (F, L). HIS-tag: red; DAPI: Blue. Arrow points to the epithelial cells with FOXL2 overexpression. N=6.

**Suppl. Fig. 3**. FOXL2$^{oe}$ transcriptome was positively correlated with transcriptome of multiple human endometriosis data sets. Positive correlation of FOXL2$^{oe}$ transcriptome with the transcriptome of ectopic endometrium from patients with severe endometriosis compared to the eutopic tissues (GSE5108 [34], A), the transcriptome of ectopic endometrium compared to normal endometrium (GSE87809 [36], B) the transcriptome of endometrium with mildly severe endometriosis compared to normal endometrium (GSE51981 [5], C), the transcriptomic of cultured stromal cells from ectopic compared to eutopic endometriosis tissues (GSE47361 [37], D). The number of up- and down-regulated genes in each dataset were labeled accordingly near the up and down arrows. The number of overlapped genes between both datasets were plotted in the venn diagram. The significance of the overlap was showed below the venn diagram. Furthermore, the overlapping genes between two datasets were divided into four groups based on its regulation direction up or down in either dataset. The significance and number of overlapped genes in these four groups were displayed in the bar graph.

**Suppl. Fig. 4**. *Foxl2* transgene expression was not detected in the diestrus ovary but found in the super-ovulated ovary of the FOXL2$^{oe}$ mice. FOXL2 was highly expressed in the small and medium follicles of diestrus ovaries from *Pgr*$^{cre}$ (A) and FOXL2$^{oe}$ (B) mice. HIS-tag staining showed no *Foxl2* transgene expression in the diestrus ovaries of *Pgr*$^{cre}$ (C) and FOXL2$^{oe}$ (D) mice. 16h after hCG injections, *Foxl2* transgene detected by His-tag was found in peri-ovulatory follicles and corpus luteum of the ovaries from FOXL2$^{oe}$ (F, H) mice, but not from *Pgr*$^{cre}$ (E, G) mice. The number of collected oocytes from the oviduct after superovulation were decreased in the FOXL2$^{oe}$ ovaries (I). Since all the ovaries showed corpus lutea after superovulation, the percentage of mice with corpus luteum were 100% for both *Pgr*$^{cre}$ (A) and FOXL2$^{oe}$ mice (J). Arrow indicates cells with positive staining of HIS-tag. N=6.

**Suppl. Fig. 5**. Hormone levels of *Pgr*$^{cre}$ and FOXL2$^{oe}$ mice. The serum levels of progesterone (A), 17β-estradiol (B), FSH (C) and LH (D) were comparable between 3M *Pgr*$^{cre}$ and FOXL2$^{oe}$ mice at diestrus stage. 16h after hCG injections, progesterone levels were increased in both *Pgr*$^{cre}$ and FOXL2$^{oe}$ mice to the similar levels (E). Each dot indicates one individual sample. Dots with solid fill represents detected reads, and dots without fill are below the detectable ranges, and showed as the lowest range. FSH, Follicle stimulating hormone; LH, luteinizing hormone; M: months old. N=6.

Suppl. Fig. 1

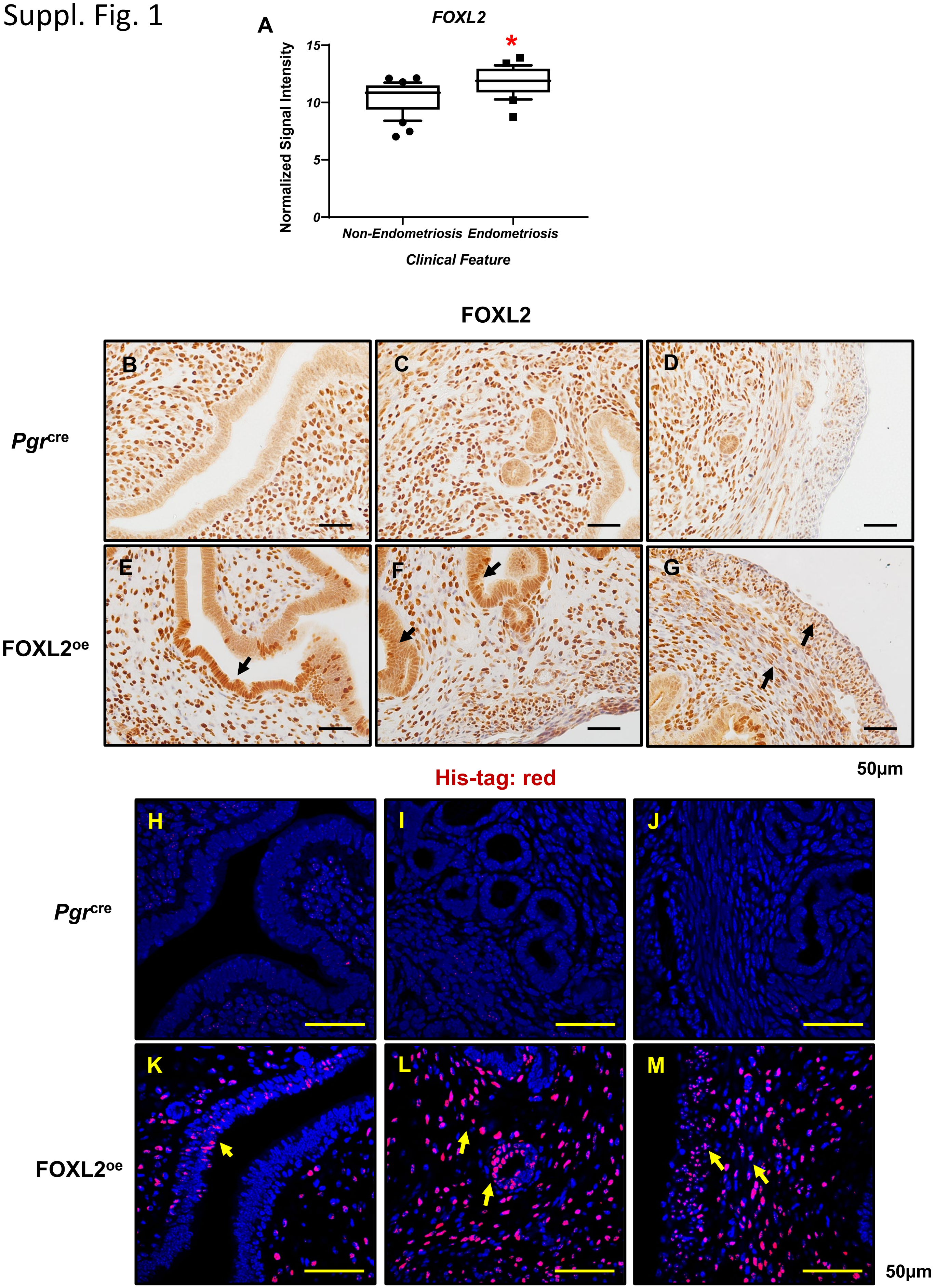
A
FOXL2
*
15
10
5
0
Normalized Signal Intensity
Non-Endometriosis
Endometriosis
Clinical Feature
FOXL2
B
C
D
E
F
G
Pgrcre
FOXL2oe
50µm
His-tag: red
H
I
J
K
L
M
Pgrcre
FOXL2oe
50µm

Suppl. Fig. 2

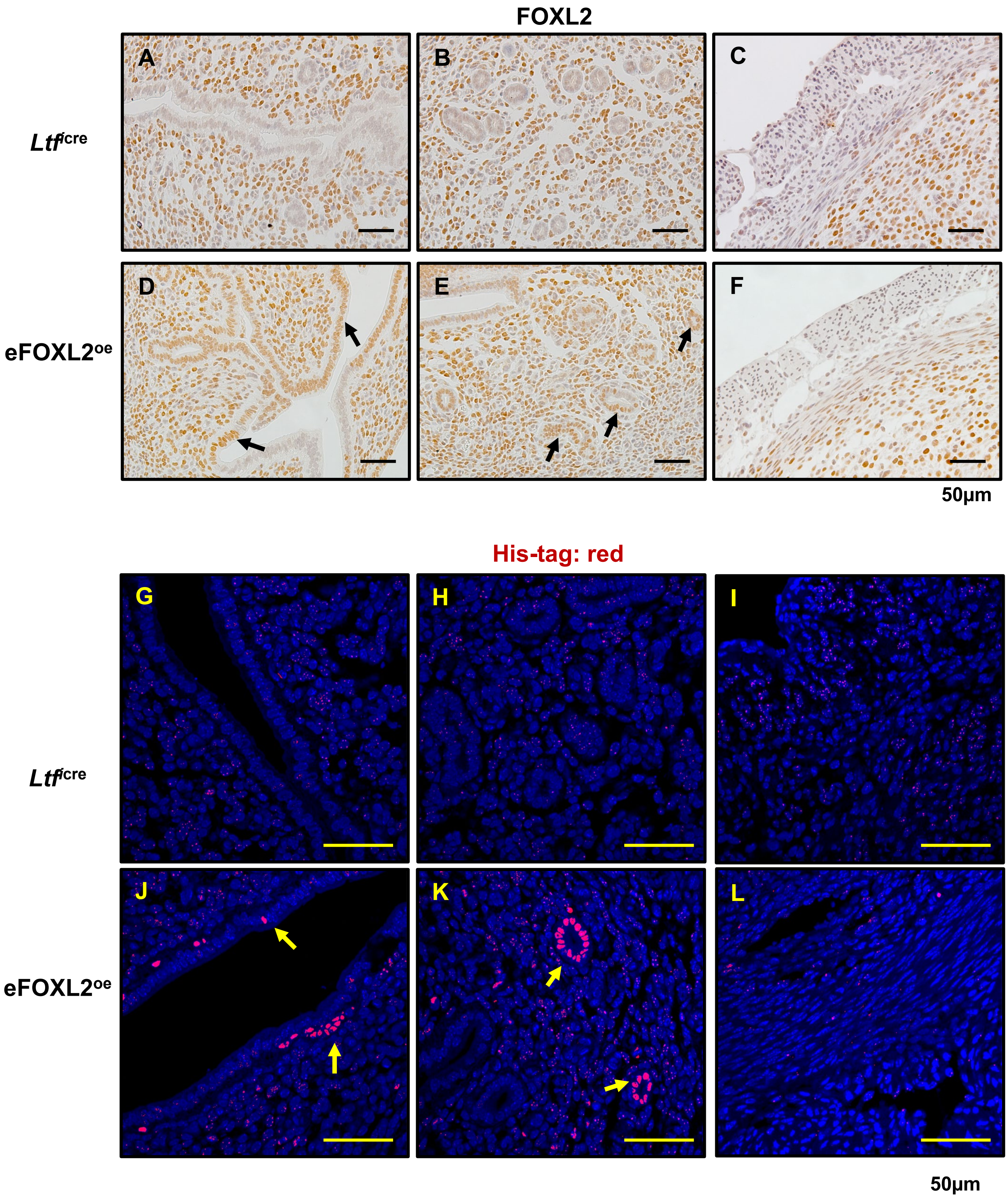
FOXL2
A
B
C
D
E
F
*Ltf*icre
eFOXL2oe
50µm
His-tag: red
G
H
I
J
K
L
*Ltf*icre
eFOXL2oe
50µm

Suppl. Fig. 3

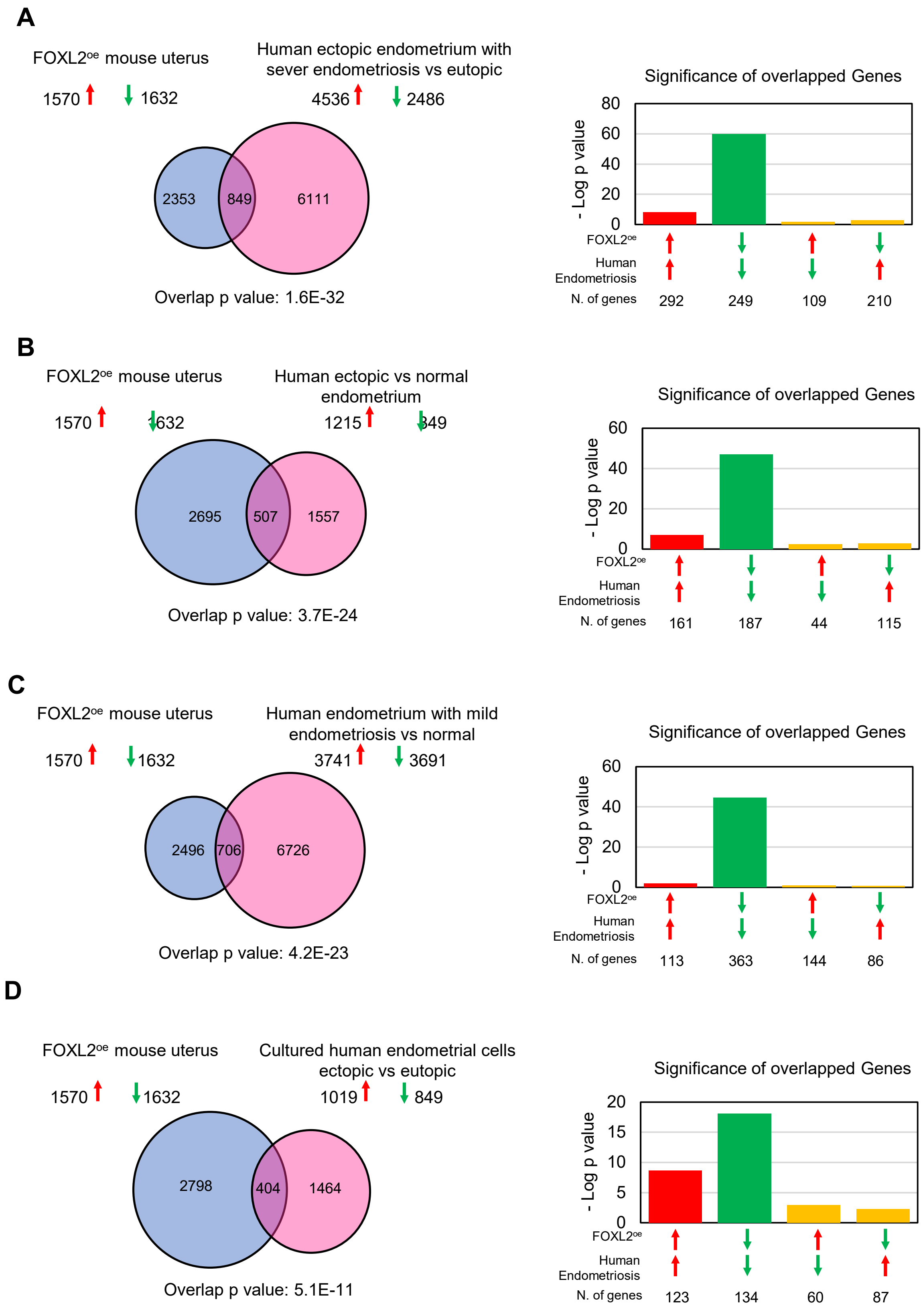

A
FOXL2oe mouse uterus
1570
1632
Human ectopic endometrium with sever endometriosis vs eutopic
4536
2486
2353
849
6111
Overlap p value: 1.6E-32
Significance of overlapped Genes
- Log p value
80
60
40
20
0
FOXL2oe
Human Endometriosis
N. of genes
292
249
109
210
B
FOXL2oe mouse uterus
1570
1632
Human ectopic vs normal endometrium
1215
849
2695
507
1557
Overlap p value: 3.7E-24
Significance of overlapped Genes
- Log p value
60
40
20
0
FOXL2oe
Human Endometriosis
N. of genes
161
187
44
115
C
FOXL2oe mouse uterus
1570
1632
Human endometrium with mild endometriosis vs normal
3741
3691
2496
706
6726
Overlap p value: 4.2E-23
Significance of overlapped Genes
- Log p value
60
40
20
0
FOXL2oe
Human Endometriosis
N. of genes
113
363
144
86
D
FOXL2oe mouse uterus
1570
1632
Cultured human endometrial cells ectopic vs eutopic
1019
849
2798
404
1464
Overlap p value: 5.1E-11
Significance of overlapped Genes
- Log p value
20
15
10
5
0
FOXL2oe
Human Endometriosis
N. of genes
123
134
60
87

Suppl. Fig. 4

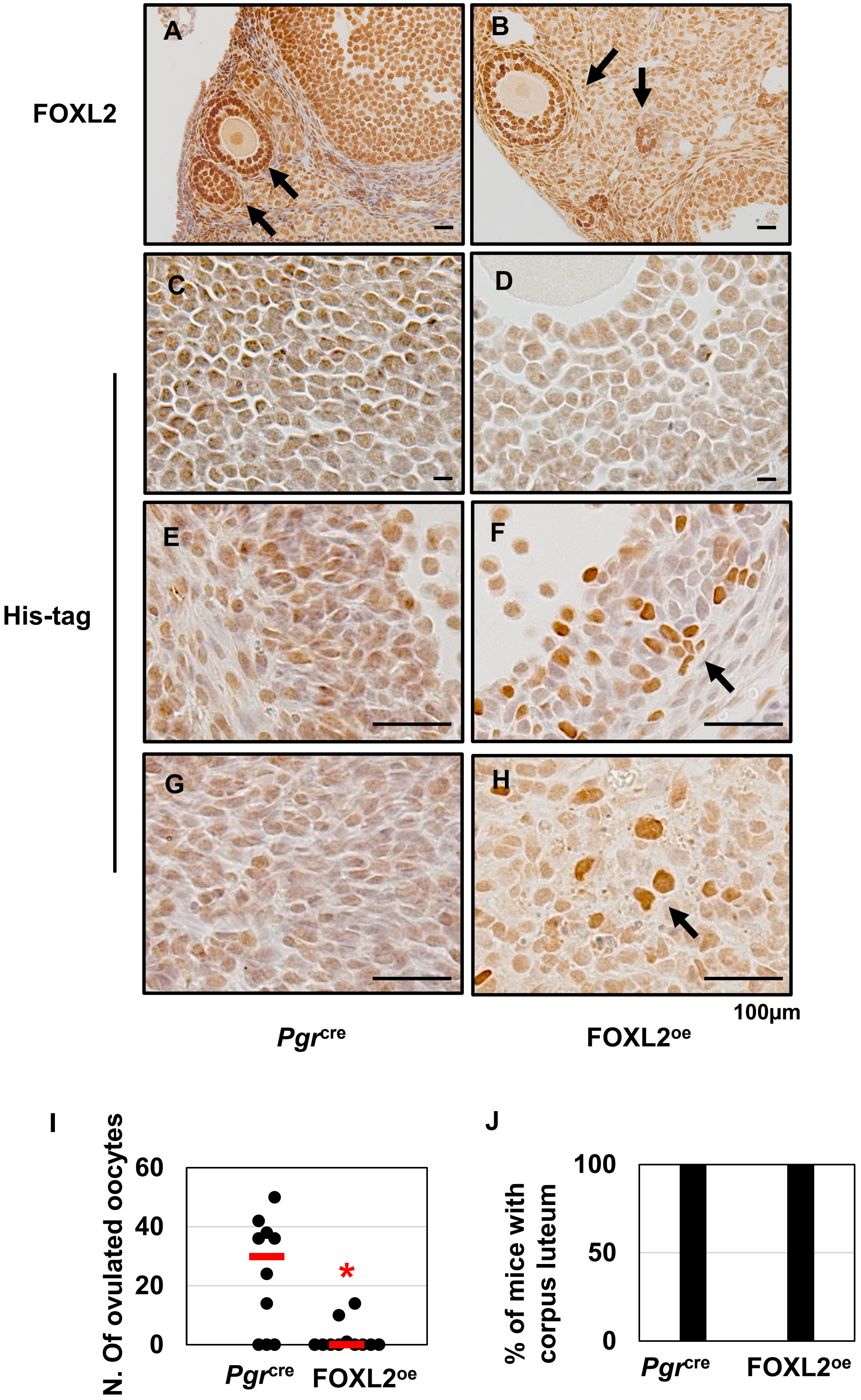

FOXL2
His-tag
A
B
C
D
E
F
G
H
100µm
Pgr^cre
FOXL2^oe
I
N. Of ovulated oocytes
60
40
20
0
*
Pgr^cre
FOXL2^oe
J
% of mice with corpus luteum
100
50
0
Pgr^cre
FOXL2^oe

Suppl. Fig. 5

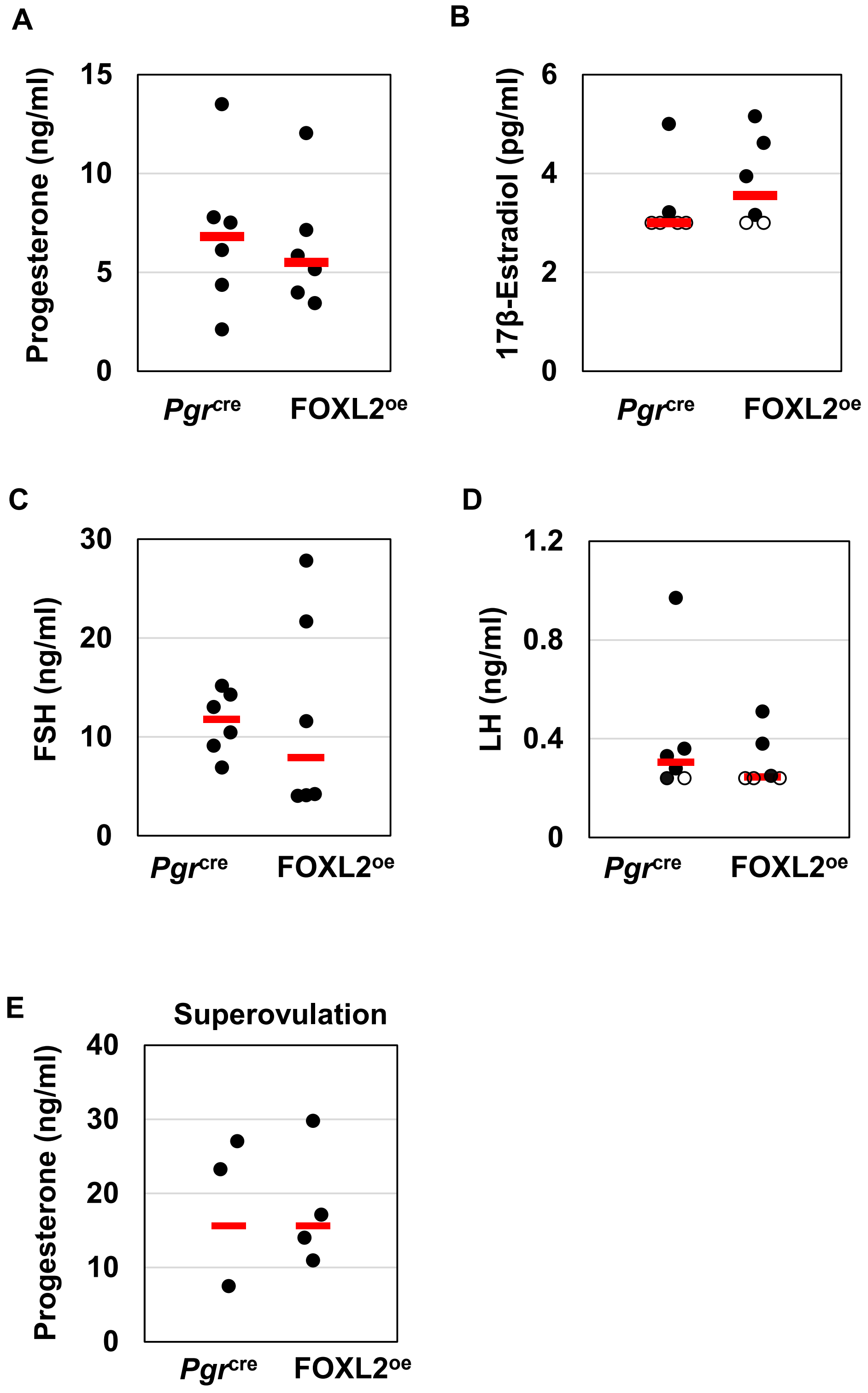

A
Progesterone (ng/ml)
15
10
5
0
Pgr^cre
FOXL2^oe
B
17β-Estradiol (pg/ml)
6
4
2
0
Pgr^cre
FOXL2^oe
C
FSH (ng/ml)
30
20
10
0
Pgr^cre
FOXL2^oe
D
LH (ng/ml)
1.2
0.8
0.4
0
Pgr^cre
FOXL2^oe
E
Superovulation
Progesterone (ng/ml)
40
30
20
10
0
Pgr^cre
FOXL2^oe
